# LLIN Evaluation in Uganda Project (LLINEUP) – The durability of long-lasting insecticidal nets treated with and without piperonyl butoxide (PBO) in Uganda

**DOI:** 10.1101/2022.02.17.480046

**Authors:** Frank Mechan, Agaba Katureebe, Violet Tuhaise, Martin Mugote, Ambrose Oruni, Ismail Onyige, Kawesa Bumali, Jonathan Thornton, Kilama Maxwell, Mary Kyohere, Moses R Kamya, Peter Mutungi, Simon P Kigozi, Adoke Yeka, Jimmy Opigo, Catherine Maiteki-Sebuguzi, Samuel Gonahasa, Janet Hemingway, Grant Dorsey, Lisa J Reimer, Sarah G Staedke, Martin J Donnelly, Amy Lynd

**Author notes:** Equal contributors (Frank Mechan, Agaba Katureebe). **Corresponding author:** Amy Lynd, Department of Vector Biology, Liverpool School of Tropical Medicine, UK.

## Abstract

Long Lasting Insecticidal Nets (LLINs) supplemented with the synergist piperonyl butoxide have been developed in response to growing pyrethroid resistance however their durability in the field remains poorly described. A pragmatic cluster-randomised trial was embedded into Uganda’s 2017-2018 distribution to compare the durability of LLINs with and without PBO

A total of 104 clusters were included with each receiving one of four LLIN products, two with pyrethroid+PBO (Olyset Plus and PermaNet 3.0) and two pyrethroid-only (Olyset Net and PermaNet 2.0). Nets were sampled at baseline, 12, and 25 months post-distribution to assess physical condition, chemical content, and bioefficacy. Physical condition was quantified using proportionate Hole Index and chemical content measured using high-performance liquid chromatography. Bioefficacy was assessed with three-minute WHO Cone and Wireball assays using pyrethroid-resistant *An. gambiae*, with 1hr knockdown and 24hr mortality recorded.

There was no difference in physical durability between LLIN products assessed (p=0.644). The pyrethroid content of all products remained relatively stable across timepoints but PBO content declined by 55% (p<0.001) and 58% (p<0.001) for Olyset Plus and PermaNet 3.0 respectively. Both PBO LLINs were highly effective against pyrethroid-resistant mosquitoes when new, knocking down all mosquitoes. However, bioefficacy declined over time with Olyset Plus knocking down 45.72% (95% CI: 22.84-68.62, p=0.021) and Permanet 3.0 knocking down 78.57% (95% CI: 63.57-93.58, p<0.001) after 25 months.

Here we demonstrate that both Olyset Plus and PermaNet 3.0 are as durable as their pyrethroid-only equivalents and had superior bioefficacy against pyrethroid-resistant *An. gambiae*. However, the superiority of PBO-LLINs decreased with operational use, correlating with a reduction in total PBO content. This decline in bioefficacy after just two years is concerning and there is an urgent need to assess the durability of PBO LLINs in other settings.

## INTRODUCTION

Long Lasting Insecticidal Nets (LLINs) are the cornerstone of global malaria control strategies, forming a physical and chemical barrier against the bites of *Anopheles* mosquitoes^(1–3)^. Progress in reducing malaria burden in sub-Saharan Africa achieved in the first decade of the 21^st^ century has been attributed, in large part, to mass distribution of LLINs^(3)^. LLINs are intended to maintain an effective level of protection for at least three years, with the expectation that distributions will take place at two-to-three-year intervals^(4, 5)^. However, recent studies suggest that the lifespan of LLINs may be less than three years^(6–8)^. To ensure the continued success of malaria control efforts, National Malaria Control Programmes (NMCPs) must identify LLIN products that demonstrate durability within the socio-economic and environmental context of their country.

The WHO currently only recommends the use of pyrethroid and pyrrole insecticides on LLINs^(9)^ and the effectiveness of LLINs is threatened by widespread pyrethroid resistance^(1, 10, 11)^. The development of target site alterations and metabolic resistance enables mosquitoes to better tolerate insecticide exposure, increasing the probability they will obtain a bloodmeal and survive the encounter^(12–14)^. While there is evidence that pyrethroid LLINs retain some protective effect against resistant mosquito populations ^(15, 16)^, the threat of resistance has incentivised the development of new classes of LLIN. Due to the limited alternatives to pyrethroids, initial efforts to maintain the impact of LLINs have focused on secondary compounds that restore the susceptibility of pyrethroid-resistant mosquitoes. Piperonyl butoxide (PBO) is a synergist that inhibits the cytochrome P450 enzymes within the mosquito which can detoxify insecticides ^[19,20]^. In 2017, the WHO provided an interim endorsement of use of pyrethroid LLINs containing PBO in areas of moderate pyrethroid resistance^[21]^ and a 2021 Cochrane review concluded that in areas of high pyrethroid resistance, PBO-LLINs nets epidemiological efficacy and were associated with a sustained reduction in parasite prevalence^[9]^. However, the same review emphasised that evidence of the durability of these PBO-LLINs under operational conditions is lacking.

LLINs are known to lose insecticide content during routine use^(17)^. As nets are handled and washed, the insecticide at the surface is depleted then gradually regenerated by a reservoir within the fibres^(18)^. Pyrethroid LLINs are designed with sufficient insecticide reserves to continue regenerating for at least three years, with the expectation they will be replaced before this time^(4)^. Currently, WHO LLIN durability guidelines quantify performance against objective bioefficacy benchmarks to assess if a three-year operational lifespan is achieved^(19, 20)^ and there is emerging evidence to suggest that bioefficacy varies substantially between products and may fall below defined efficacy thresholds within three years^(6–8)^.

In Uganda, the country with the highest malaria burden in East Africa, progress in controlling transmission has faltered^[28]^. The declining efficacy of conventional control strategies coincides with emerging evidence of both high levels of knockdown resistance (*kdr*) and metabolic resistance in mosquito populations throughout the country^(21, 22)^. As part of a commitment to achieve universal coverage of LLINs, the Ugandan Ministry of Health initiated a mass distribution of LLINs and PBO LLINs in 2017. A randomised control trial was embedded within this distribution programme to evaluate the impact of LLINs with and without PBO^(23)^. From this, it was demonstrated that PBO-LLINs reduce parasite prevalence in children aged 2-10 years old and vector density more effectively than conventional LLINs for at least 25 months^(24, 25)^. The current study was conducted as part of the same trial to evaluate the durability of the PBO-LLINs. Here the physical integrity, chemical integrity, and bioefficacy of two PBO-LLIN products are assessed in comparison with their pyrethroid-only equivalents at 12 and 25 months post distribution.

## METHODS

### Study site

The trial protocol for this study has been published^(23).^ A total of 104 clusters (health sub-districts) in eastern and western Uganda were randomly assigned to receive one of four LLIN products, including two LLINs with PBO (PermaNet 3.0 and Olyset Plus) and two LLINs without PBO (PermaNet 2.0 and Olyset Net). Cross-sectional community surveys were carried out in 50 households per cluster (5200 households per survey) to confirm the presence of the expected LLIN product from the distribution and entomological surveillance undertaken in 10 households per cluster. Efficacy data from this study have been published previously^(24)^. In the current study, we quantify the chemical and physical integrity of 400 LLINs, 97 −100 nets of each type (Supplementary Table 1), withdrawn from households after 12 months and 25 months (800 nets total). These nets were assessed alongside unused nets of the same LLIN products.

### LLIN description

Four LLIN products were distributed and assessed in this study; Olyset Net treated with permethrin, PermaNet 2.0 treated with deltamethrin, Olyset Plus treated with permethrin and PBO, and PermaNet 3.0 treated with deltamethrin and incorporating PBO on the top surface of the net only. All nets were 180cm long × 170cm wide × 170 high; the chemical and fabric specifications of each LLIN product are shown in Table 1.

**Table 1.**
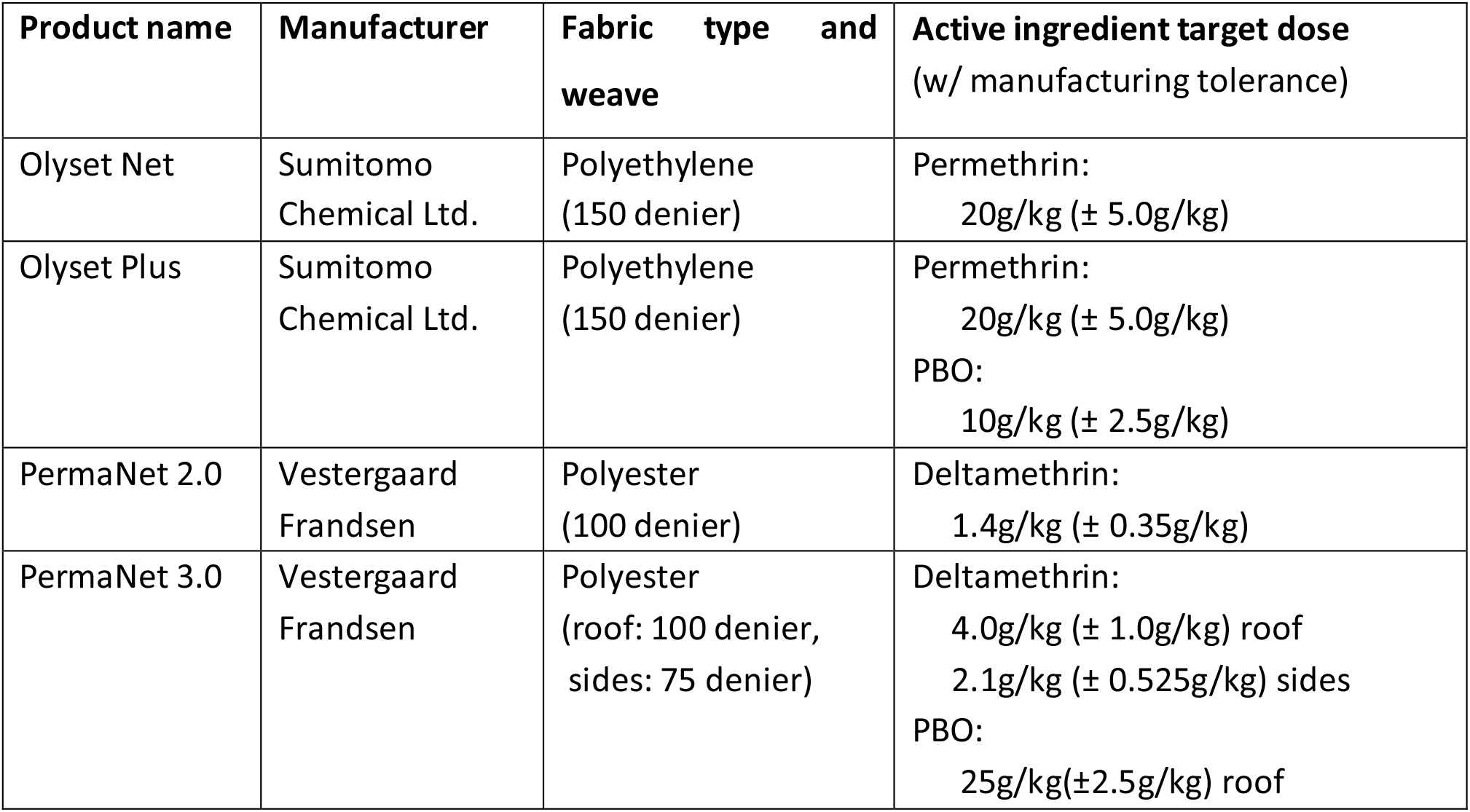
Specifications of LLIN products assessed in study. The target dose was defined as the amount of chemical per kg of fabric

### Field collections

Net sampling was performed at baseline, 12 months, and 25 months post-distribution. At baseline, a total of 20 nets were retained (five of each LLIN product) from the LLINs that were to be distributed during the campaign to be used as baseline samples. Post-distribution, at 12 and 25 months, 100 LLINs of each type were collected from houses enrolled in the community survey.

Nets were sampled with replacement, exchanged for a new net of the same type. If no study net was found at the selected household or the net was an unexpected type then the next household on the reserve list was sampled instead. No more than one net per household was sampled. Information on the construction of the dwelling was recorded, with the household categorised as ‘improved’ if it had both brick walls and an iron roof. Otherwise, the dwelling was categorised as ‘traditional’.

On collection, sampled nets were labelled and placed individually in zip-lock bags. All sampled nets were transported to the project office in Bugembe, Jinja for physical assessment and processing. After physical measurements were recorded, seven 30cmx30cm pieces were cut from each net (one from each side panel and three from the top) and samples sent to the Liverpool School of Tropical Medicine (Liverpool. United Kingdom) for chemical and bioefficacy assessment.

### Physical integrity

To assess the physical integrity of the net fabric, nets were placed over a metal frame measuring W160 × L180 × H170 cm and any holes >0.5cm recorded^(26)^. The size of a hole was defined by its length (the longest dimension) and width (measurement perpendicular to length measurement). Holes smaller than 0.5cm (in length or width) and holes that had been repaired were noted but not included in the final dataset. Hole size was calculated using the formula for an ellipse (area=π*length*width). The total area of damage on a net was summed and used to categorise the net within the WHO proportionate Hole Index (pHI) categories; ‘Good’ (0-64cm^2^), Damaged’ (65-642cm^2^), or ‘Too torn’ (643cm^2^+)^(17)^. Additionally, the proportion of nets of each LLIN product was calculated for each timepoint.

The vertical height of the hole on the net surface was measured as the distance between the ground and the centroid of the hole (no location measurements were taken for holes on the top surface). This height value was used to locate the hole within Top, Upper, and Lower sectors of an LLIN as defined by Sutcliffe and Yin^(27, 28)^. Following physical integrity testing, two 30cm × 30cm square net pieces were sampled from the top of each LLIN for bioefficacy and chemical assessment. The samples were wrapped in aluminium foil and stored at room temperature prior to use in WHO cone bioassays. Samples were subsequently stored at 4°C until chemical integrity could be assessed.

### Chemical integrity

To quantify the content of active ingredients, chemical analysis was performed using high-performance liquid chromatography (HPLC) after extraction in 10% 1-propanol in heptane. A total of 30 nets of each LLIN type were analysed at each timepoint, with two samples taken from each net.

The HPLC analysis was performed on an Agilent 1100 Series machine (Aglient, California US) at a wavelength of 226nm, using a modification of the methods outlined by Ngufor *et al.*, 2022 ^(29)^. Quantities of Permethrin, Deltamethrin, and Piperonyl Butoxide were calculated by comparison to standard curves of each compound (PESTANAL®, analytical standard, Sigma-Aldrich, Missouri US) and corrected against internal standard dicyclohexyl phthalate (DCP). HPLC data were analysed using OpenLAB software v2.1 (Aglient, California US).

### WHO cone bioassays

To assess bioefficacy, WHO cone bioassays were performed using the protocol outlined in WHO durability monitoring guidelines^(19, 20)^.

All bioefficacy testing was performed on nets also assessed in chemical analysis. The two pieces from each net were each tested in duplicate, thus a total of four cone exposures were performed per net. Cone bioassay design followed WHO protocol but with the testing board angled at 45 degrees^(30)^. Ambient conditions in the testing room were targeted to 27°C ± 2°C and 80% ± 10% relative humidity. All mosquitoes used were 3-5-day old unfed females, reared in temperature and humidity-controlled insectaries. Approximately 25 mosquitoes were used in each cone exposure assay per net piece for each mosquito strain.

Two different mosquito strains were used in the cone bioassays: ‘Kisumu’ and ‘Busia’. Kisumu is a pyrethroid susceptible strain of *An. gambiae* collected in 1975 from what is now Kisumu county (formerly Kisumu district), in Western Kenya. Busia is a strain established in November 2018 from mosquitoes collected in Busia, Eastern Uganda by Ambrose Oruni. Busia has been previously characterised as possessing resistance to pyrethroids through both target site alterations (*Vgsc*-1014S) and metabolic resistance mechanisms (*Cyp4j5, Cyp6aa1* and *Coeae1d*)^(21, 22).^ WHO tube assays with standard discriminating doses indicate Busia is more resistant to permethrin than deltamethrin (Supplementary Figure 1).

WHO bioefficacy criteria is defined as the proportion of nets that achieve either 80% mortality or 95% knockdown against pyrethroid susceptible *An. gambiae s.s.* mosquitoes. An LLIN product was considered to have passed if 80% of nets met these criteria at all timepoints up to 24 months. Chemical and physical integrity data are not included in bioefficacy criteria.

### WHO wireball assays

Due to previous literature indicating that WHO cones tests are insufficient to assess the bioefficacy of LLIN products containing insecticides with high contact irritancy^(31, 32)^, such as permethrin, supplemental WHO wireball assays were performed on the same samples used in WHO cone bioassays. In the WHO wireball method, the net to be tested is affixed around a wire cube measuring 15cmx15cmx15cm. As in the cone bioassay, seven 3-5 day old females were released into the wireball for three minutes then assessed for 1hr knockdown and 24 hour mortality.

### Data analysis

Data analyses were conducted using R (version 3.6.0), all graphs were produced using the ggplot2 package (version 3.2.1). Associations between outcomes and variables of interest were quantified using Generalized Linear Mixed Models (GLMMs) using the ‘lme4’ package (version 1.1-21). To account for unexplained variation between separate pieces from individual nets and between clusters, the net ID (a unique identifier for each net distributed) and HSD number were each included in the models as a random effect. The model selection process used stepwise regression, working backwards from a maximally complex model to produce the most parsimonious fit. Variables that did not significantly increase explanatory power, as indicated by log-likelihood ratio tests (LRTs)(‘lmtest’ package version 0.9-37), were excluded from the final model. The p values reported are the output of these LRTs. Pairwise comparisons between levels within a categorical variable were performed using least square means with the ‘lsmeans’ package (version 2.30-0).

To quantify the relationship between chemical integrity and bioefficacy, the HPLC outputs for each net were combined with their corresponding WHO cone assay or WHO wireball assay mortality data (for PermaNet 3.0 and Olyset Plus respectively). A GLMM was then fit separately to the PermaNet 3.0 and Olyset Plus data, with pyrethroid content and PBO content each fit as a fixed effect. Model selection and p value reporting was performed as above. The 3D plots were produced using the ‘plot3D’ package (version 1.4).

### Ethical approval

The trial was approved by the Ugandan National Council for Science and Technology (UNCST Ref HS 2176), Makerere University School of Medicine Research & Ethics Committee (SOMREC 2016-133), London School of Hygiene & Tropical Medicine Ethics Committee (LSHTM Ref 12019), and the Liverpool School of Tropical Medicine (LSTM Ref 16-072), which is the sponsoring institute.

## RESULTS

### Physical integrity

#### Proportion of nets in each pHI category

At 12 months post-distribution, the proportion of nets classified as ‘too torn’ on the pHI scale was 0.066 (Fig 1A), with this proportion approximately doubling after 25 months (Fig1B) to 0.125 (OR: 2.017, 95%CI: 1.268-3.208, p<0.001, Supplementary Table 2). There was no significant difference in the proportion of nets that were ‘too torn’ between LLIN products (p= 0.644).

**Fig 1.**
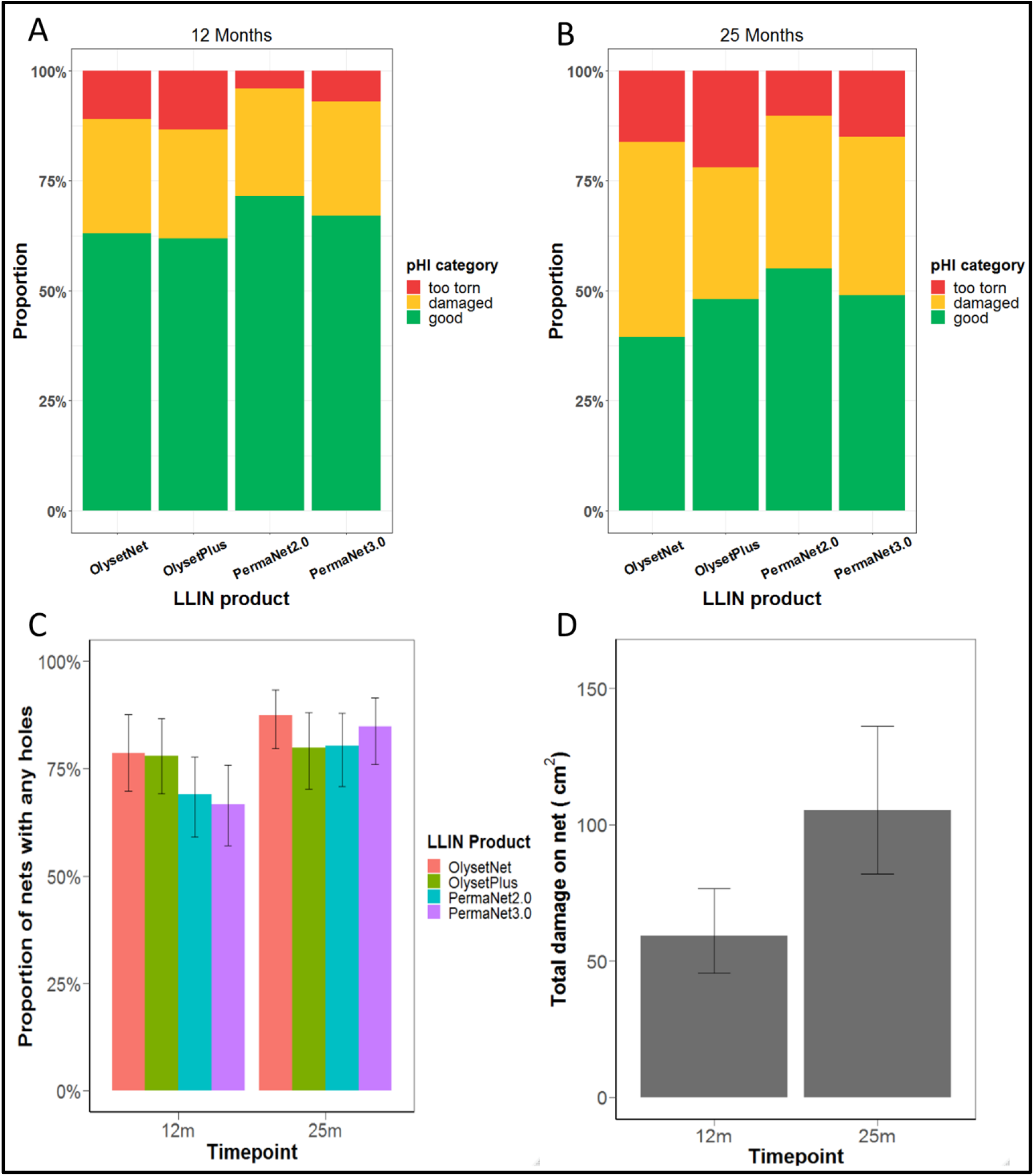
Physical integrity outcomes at 12- and 25-months post-distribution. **A)** Proportion of collected nets in each pHI category (‘Too torn’, ‘Damaged’, ‘Good’) at 12 months. **B)** Proportion of collected nets in each pHI category at 25 months. **C)** Proportion of nets with at least one hole. **D)** Mean total surface area of damage per net at 12 and 25 months post-distribution across all LLIN products.

When categorised by the type of housing they were collected from, it was observed that nets from traditional housing were more likely to be in poor physical condition than those from improved housing (OR: 3.350, 95%CI: 1.865-6.016, p=0.003, Supplementary Table 2). After 25 months in operational use, the proportion of nets from traditional housing categorised as ‘too torn’ was 0.297 compared to 0.112 for improved housing (Supplementary Figure 2).

#### Proportion of nets with at least one hole

The proportion of nets of each type with at least one hole at 12- and 25 months post distribution is shown in Figure 1C. The overall proportion of nets with at least one hole after 12 months in operational conditions was 0.727, increasing to 0.829 after 25 months (OR: 1.821, 95%CI: 1.289-2.571, p<0.001). There was no difference in the proportion of nets with at least one hole between the four LLIN products tested at any timepoint (p=0.306).

#### Total surface area of holes

There was no difference in total hole area between any of the four LLIN products tested (p=0.270). However, across all net types there was an overall increase in holed area from 12m post-distribution to 25m post-distribution (p=0.0005, Fig 1D), which approximately doubled from 59.33cm^2^ (95% CI: 45.08 – 78.25) to 105.49cm^2^ (95% CI: 83.43 −136.86).

### Chemical integrity

At baseline, all net samples tested met or exceeded the minimum target dose of active ingredients per their respective manufacturer specifications.

#### Deltamethrin

The deltamethrin content of PermaNet 3.0 was lower at each subsequent timepoint (p=<0.001, Fig 2A). In the period from baseline to 25 months, mean deltamethrin content of PermaNet 3.0 nets declined from 4.98g/kg (95% CI: 4.08-6.01) to 3.484g/kg (95% CI: 3.19–3.78). Despite this, the deltamethrin content of all PermaNet 3.0 nets collected at 25 months remained within the range of the target dose (3.0-5.0g/kg). For PermaNet 2.0, mean deltamethrin content after 25 months was not statistically different from baseline (p=0.071).

**Fig. 2.**
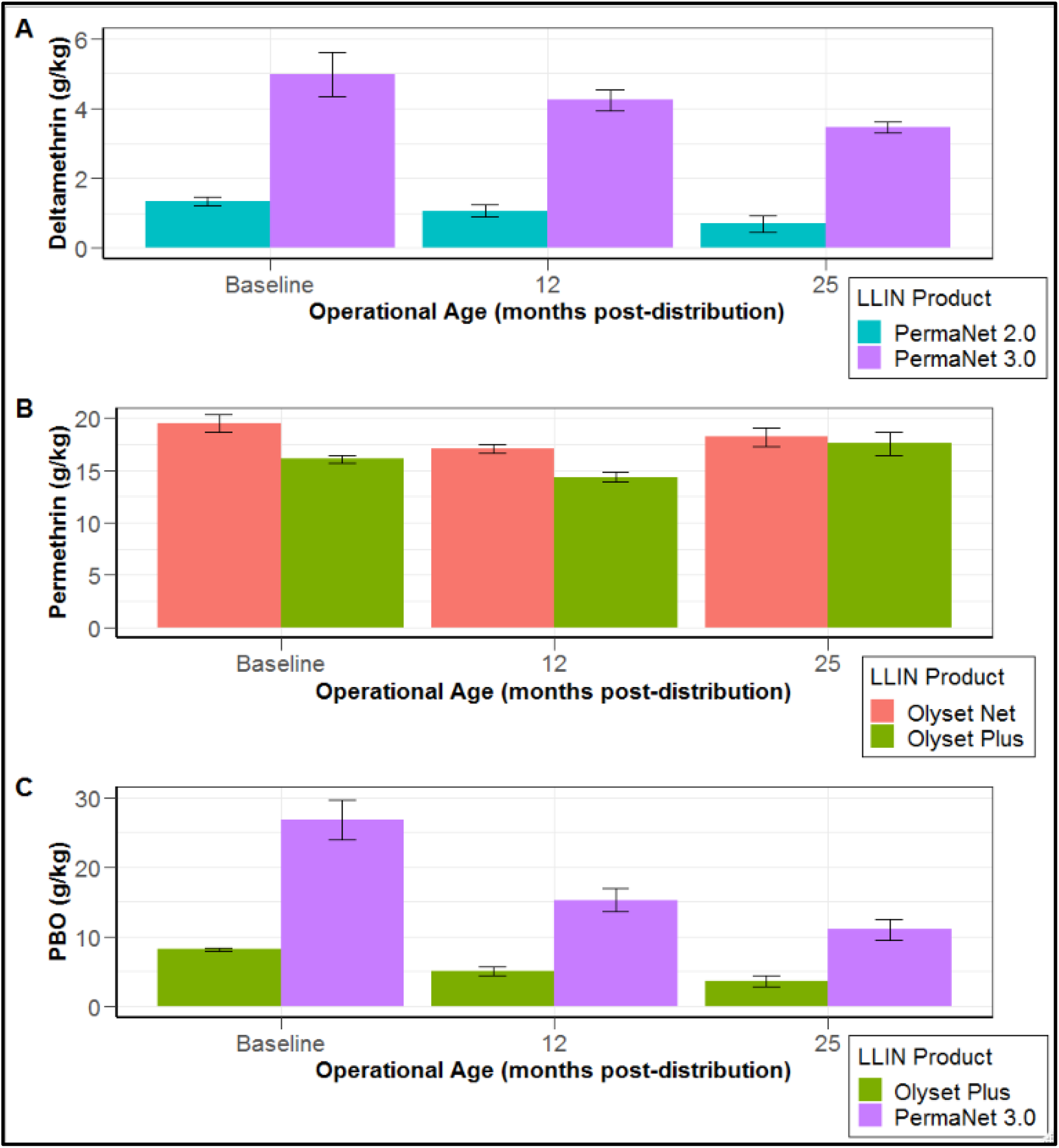
Mean concentration of **(A)** deltamethrin, **(B)** permethrin, and **(C)** PBO detected in net samples at each sampled timepoint (measured using HPLC). Error bars indicate 95% CIs.

#### Permethrin

The permethrin content of Olyset Plus varied across the sampled timepoints (p<0.001, Fig 2B) however pairwise comparison indicated no overall difference between baseline and the final timepoint at 25 months (p=0.591). Mean permethrin content in Olyset Plus at baseline was 16.08 (95% CI: 13.70-18.62), declining to 14.54 (95% CI: 13.64-15.35) after 12 months, then increasing to 17.39 (95% CI: 16.53-18.22) after 25 months. A similar pattern was observed for Olyset Net, with permethrin content varying across timepoints overall (p<0.001) yet pairwise comparison indicating no overall difference between baseline and the 25-month timepoint (p=0.327).

#### PBO

The PBO content of PermaNet 3.0 declined across the sampled timepoints (p<0.001, Figure 2C). PBO content for PermaNet 3.0 at baseline was 26.81g/kg (95% CI: 22.80-31.07) before declining sharply to 15.28g/kg (95% CI: 13.74-16.71) after 12 months (p=0.001), then falling further to 11.03g/kg (95% CI: 9.35-12.67) after 25 months (p=0.001).

A similar downwards trend in PBO was observed for Olyset Plus across timepoints (p<0.001). At baseline mean PBO content was 8.17g/kg (95% CI: 6.51-9.82) before declining to 5.03g/kg (95% CI: 4.37–5.74) after 12 months (p=0.002). From 12 months to 25 months post-distribution, PBO content further fell to 3.66g/kg (95% CI: 2.97-4.28, p=0.013).

### Bioefficacy

#### Cone bioassay: Pyrethroid susceptible *An. gambiae*

All LLINs were effective per WHO definition against the pyrethroid-susceptible Kisumu strain (defined as achieving either 95% knockdown or 80% mortality), both when new and 12 months post-distribution. Overall mean cone mortality was 96.93% (95% CI: 95.77% – 98.10%) at baseline. Adjusted cone mortality was statistically indistinguishable between LLIN products (p=0.522) and did not vary significantly between timepoints (p=0.589).

#### Cone bioassay: Pyrethroid resistant *An. gambiae*

Bioefficacy against the pyrethroid-resistant strain in cone assays varied between PBO-LLINs.

Knockdown for PermaNet 3.0 remained very high throughout, achieving 99.7% (95%CI: 97.26 – 99.65, Fig. 3A) at baseline and remaining stable to 12 months (p=0.441), though declining to 78.57% (95% CI: 63.57-93.58, p<0.001) after 25 months. PermaNet 3.0 was fully lethal against the pyrethroid-resistant strain when new but mortality declined with operational use, falling by 26.8% (95%CI: 16.28-37.33) for each year in the field (p<0.001, Fig. 3B). In comparison, both mortality and knockdown with PermaNet 2.0 against the pyrethroid-resistant strain was very low at all timepoints (3% and 6% respectively).

**Figure 3.**
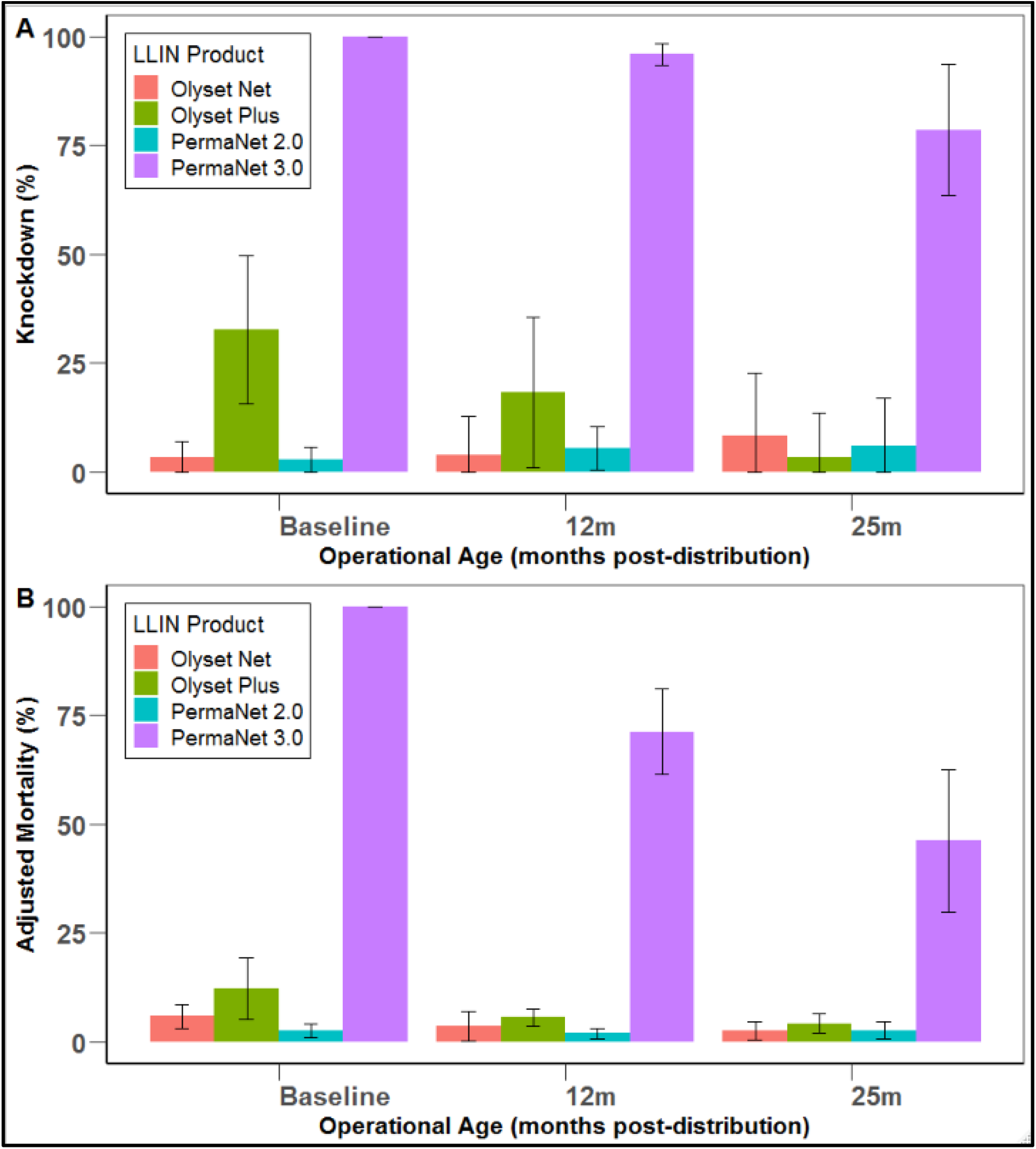
Mean **(A)** knockdown and **(B)** adjusted mortality in cone bioassays with pyrethroid resistant *An. gambiae* s.s. (‘Busia’) for each LLIN product tested at baseline, 12 months, and 25 months in the field.

Knockdown with Olyset Plus was 46.98% (95%CI: 18.55-79.13) when new but fell to considerably to 3.54% (95%CI: 0.7-10.54) after two years (p=0.005). Mortality with Olyset Plus in cone assays was low throughout, killing 12.19% (95%CI: 5.45-17.01) at baseline and 3.34% (95%CI: 0-8.71) after two years but with no significant difference between timepoints (p=0.226, Fig. 3B). Knockdown and mortality with Olyset Net was low at all timepoints (9% and 6% respectively).

#### Wireball assay: Pyrethroid resistant *An. gambiae*

The performance of Olyset Plus in wireball assays was greatly improved compared to the same nets in the cone assay, knocking down 98.93% (95%CI: 94.43-100, Fig.4A) at baseline. After 12 months knockdown had not significantly reduced (73.92%, 95% CI: 54.88-92.97, p=0.376) however there was an overall decline to 45.72% (95% CI: 22.84-68.62, p=0.021) after 25 months.

**Fig 4.**
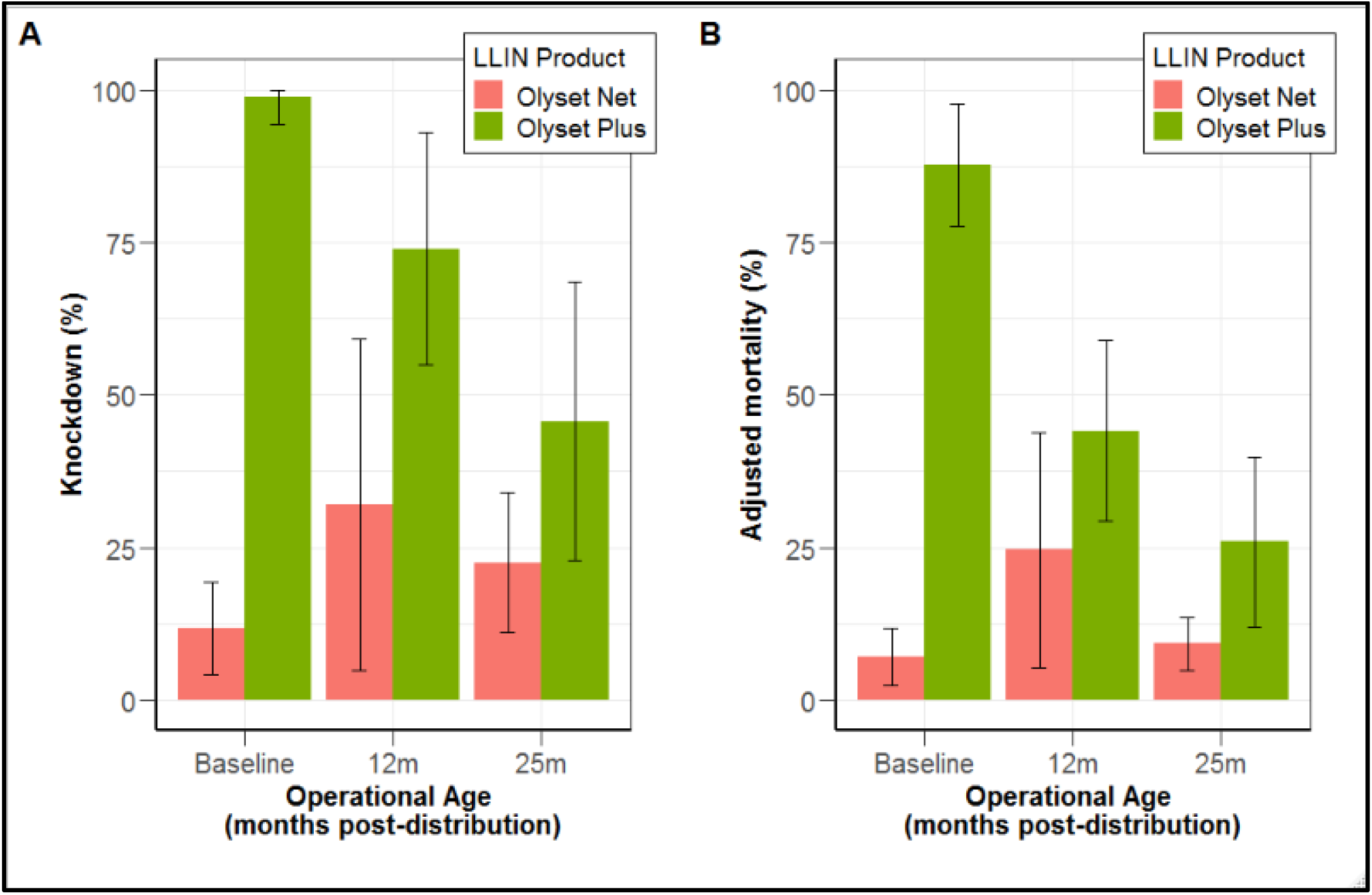
Mean **(A)** knockdown and **(B)** adjusted mortality in WHO wireball assays with pyrethroid-resistant *An. gambiae* strain (‘Busia’) for Olyset Net and Olyset Plus at baseline, 12 months, and 25 months in the field.

Mortality for Olyset Plus against the pyrethroid resistant strain in wireball assays at baseline was similarly improved compared to the cone assay, killing 87.72% at baseline (95%CI: 77.68–97.76, Fig.4B). However, after 12 months mortality had declined to 44.15% (95%CI: 29.32 – 58.98, p=0.002) though the subsequent decline to 25.92% (95%CI: 11.92-39.93) at 25 months was not statistically significant (p=0.216)

The bioefficacy of Olyset Net in the wireball assay was low at all sampled timepoints, with overall mean knockdown and mortality 22% and 13.5% respectively.

### Relationship between chemical integrity and bioefficacy

The relationship between chemical integrity and predicted mortality for the pyrethroid-resistant *An. gambiae s.s.* Busia line is shown in Figure 5.

**Figure 5.**
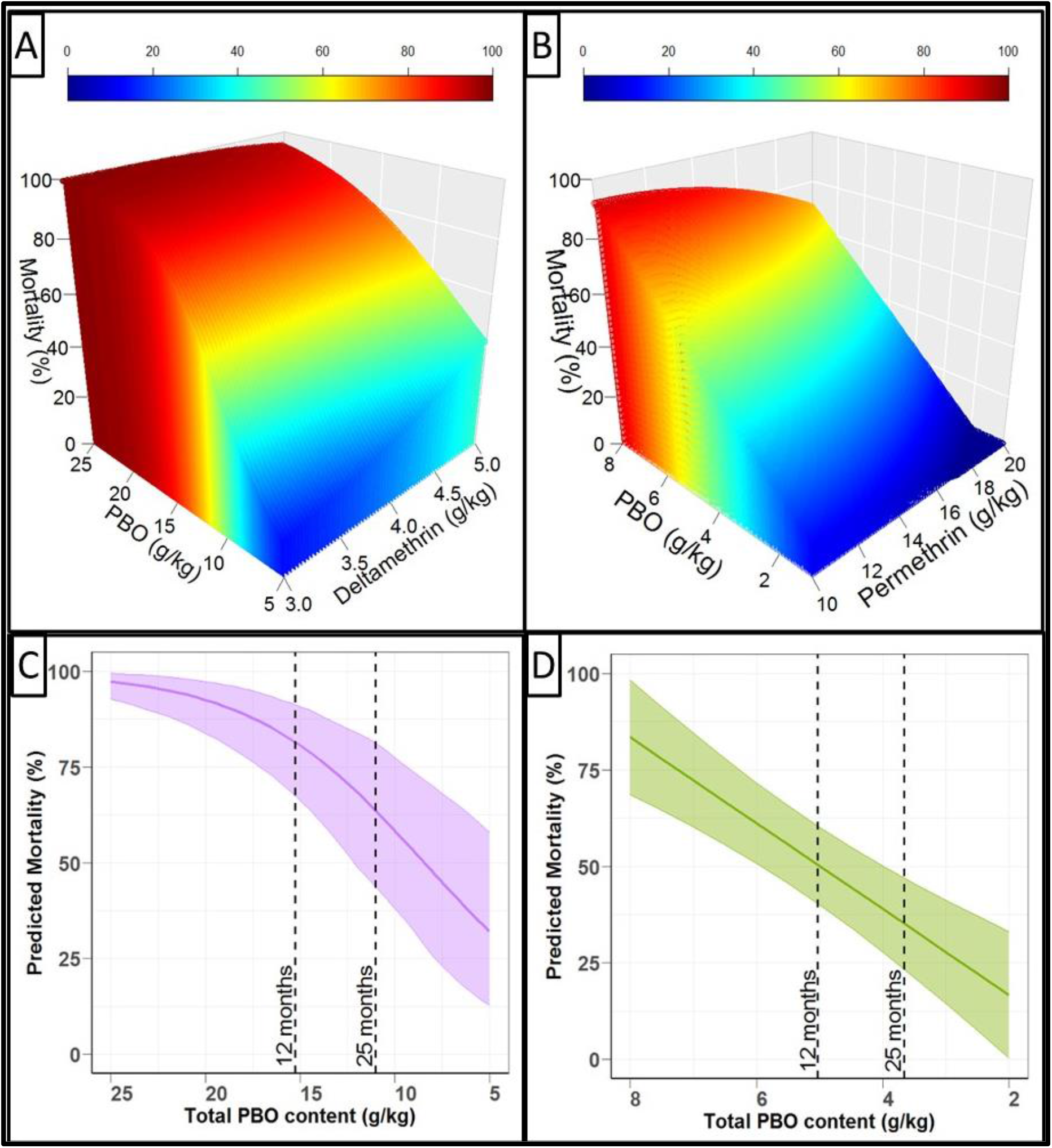
Relationship between total chemical content and bioefficacy against pyrethroid-resistant *An. gambiae s.s*. **(A)** PermaNet 3.0 in WHO Cone Bioassays **(B)** Olyset Plus in WHO Wireball Bioassays **(C)** PermaNet 3.0 in WHO Cone with deltamethrin value fixed at mean (4.42g/kg) **(D)** Olyset Plus in WHO Wireball with permethrin value fixed at mean (15.45g/kg).

For PermaNet 3.0 in the WHO cone bioassay, mortality was dependent on both total deltamethrin content and total PBO content, as indicated by a significant interaction between the two variables (p<0.001, Fig 5A). Modelling indicated there is a non-linear association between PBO content and mortality, with mortality falling more sharply with each consecutive g/kg of PBO that is lost (Fig 5C). When the deltamethrin value was fixed at the mean of the data (4.42g/kg) a reduction in PBO from 25g/kg to 15g/kg resulted in predicted mortality falling from 98% to 90%. Furthermore, a reduction in PBO content from 15g/kg to 5g/kg resulted in a decline in predicted mortality from 90% to 57%. Consequently, the model predicted that to achieve 80% mortality against this pyrethroid resistant mosquito strain a minimum of 11g/kg PBO was needed.

For Olyset Plus in the WHO wireball bioassay, mortality had no statistical relationship with total permethrin content (p=0.583) and was instead directly correlated with total PBO content (p<0.001, Fig. 5B). Modelling indicated there was a linear association between PBO content and predicted mortality, with mortality falling by 11.12% for each g/kg PBO that is lost (Fig. 5D). The model predicted that to achieve 80% mortality against this strain a minimum of 7.7g/kg PBO was needed.

## Discussion

### Physical integrity

There was no difference in physical integrity outcomes between any of the four LLINs tested after 25 months in operational conditions. Thus, PBO-LLINs nets were as physically durable as their pyrethroid-only equivalents. Furthermore, it was observed that nets sampled from ‘traditional’ thatched-roof housing were almost three times more likely to be in the most severely damaged category than nets from ‘improved’ iron-roofed housing. Consequently, there may be an argument to distribute nets more frequently than three years in regions where traditional housing remains common. However, it should be noted that the net attrition rate was high, with adequate coverage of LLINs (one LLIN for every two residents) decreasing from 71% at baseline to 35% after 25 months (Maiteki-Sebuguzi, C. *et. al.* in prep), indicating that LLIN attrition after distribution is an issue. If, as might be expected, damaged nets were discarded at a higher rate than nets in good condition, then the physical damage observed in the current study may be an underestimate.

The current physical integrity outputs outlined in the WHO durability guidelines cannot be directly interpreted in terms of personal and community protection from mosquito bites. There is a need to better understand the impact of declining physical integrity on both mosquito blood-feeding inhibition and mortality. There is empirical evidence that damage to pyrethroid LLINs reduces personal protection from bites but that mortality is independent from holed surface area and instead dependent on resistance status^(33)^. Consequently, damaged LLINs would be expected to retain some community effect against mosquito populations that are susceptible to their chemistry. Despite this, the median retention time of LLINs is well below three years in many settings (1.64 years across sub-Saharan Africa and 1.66 years for Uganda)^(34)^. Given evidence that perception of physical integrity is the primary consideration in retention^(35)^, developing more durable LLIN products may have epidemiological impacts beyond what would be indicated by studies of mosquito behaviour, due to improved retention.

Current WHO durability guidelines require the size of holes to be recorded but not their location on the net, thus all holes are weighted equally regardless of where they are. Recent behavioural experiments demonstrate that *An. gambiae* host-seeking activity is strongly focused on the top surface of the LLIN ^(27, 28, 36, 37)^. This highlights an important knowledge gap in the relationship between hole location on a net and the probability of mosquito entry and net effectiveness.

### Chemical integrity

The pyrethroid content of the LLINs assessed was relatively stable across the two years of the study, with the exception of PermaNet 3.0 which declined by ~30% (yet was still within the manufacturer’s target range). The stability of pyrethroids over two years observed here is consistent with studies from a range of settings ^[6,7,39]^. In contrast, the PBO content of both PBO-LLINs declined more rapidly over the same time period, with under half of the initial content remaining after 25 months. Nonetheless, despite this decline in PBO content the concurrent trial of epidemiological outcomes in the study site demonstrated that PBO-LLINs maintained superior protection over their conventional equivalents up to 25 months^(24, 25)^.

While a strong correlation between total PBO content and bioefficacy was observed for both PBO-LLINS, this relationship may not be causal and total chemical content quantified by HPLC may not be fully representative of the concentration at the surface bioavailable to mosquitoes^[40]^. There is currently a lack of tools for quantifying the concentration important for future studies seeking to link chemical composition to bioefficacy.

### Bioefficacy

Both Olyset Plus and PermaNet 3.0 tested demonstrated superior bioefficacy against the pyrethroid-resistant strain than their pyrethroid-only equivalents. This observation is consistent with the previously reported finding that these nets reduced childhood parasitaemia in the study area where these nets were collected^(24)^. However, while both PBO-LLINs tested were highly effective against the pyrethroid-resistant strain at baseline their bioefficacy diminished with operational use (with the mortality associated with Olyset Plus and PermaNet 3.0 decreasing to 26% and 46% respectively after two years). The diminishing differential in bioefficacy between PBO-LLINs and their pyrethroid-only equivalents is also consistent with the observation that differential impact on childhood parasitaemia narrowed over the same time. The steep reduction in bioefficacy with both PBO-LLINs against a study site specific pyrethroid-resistant strain is greatly concerning. These nets were distributed with the expectation they will be replaced after three years yet these findings indicate that they have greatly diminished killing affect after the first two years. While the bioefficacy values themselves are specific to the Busia strain, there is an urgent need to investigate if this downwards trend is observed in other settings. Given these findings, there is an argument that, within the Ugandan context, LLINs should be distributed on a two rather than three-year cycle to maintain efficacy.

The low knockdown and mortality observed with Olyset Plus in the WHO cone bioassay was in strong contrast with the high bioefficacy observed with the same nets in the WHO wireball bioassay. This difference in outcomes between methodologies may be associated with the excitorepellency of permethrin, manifesting as reduced contact with the net surface. As the wireball method surrounds the mosquito on all sides with netting, there is no insecticide-free surface to rest on and a greater insecticidal effect is observed. Consequently, while WHO guidelines were followed in the current study, future investigations with excito-repellent LLINs may wish to also include an assay that prevents avoidance from the net, such as the WHO wire-ball assay^(31)^.

## CONCLUSION

This LLIN durability study was conducted alongside a trial into the epidemiological effectiveness of PBO-LLINs in protecting against the bites of *Anopheles* mosquitoes in Uganda, where there is widespread pyrethroid resistance. Here, we demonstrate that both Olyset Plus and PermaNet 3.0 were as physically durable as their conventional equivalents and had superior bioefficacy against pyrethroid-resistant *An. gambiae s.s.* mosquitoes from the trial site. However, the superiority of PBO-LLINs over conventional LLINs in bioassays narrowed with the operational life of the net; coinciding with a decline in PBO content. Additionally, we observed that nets collected from traditional thatched-roof housing were far more likely to be severely damaged than nets from improved iron-roofed housing.

The diminished bioefficacy of PBO-LLINs against pyrethroid-resistant mosquitoes after just two years of operational use is of great concern and there is an urgent need to assess the durability of these LLIN products in other settings. Given these findings, we suggest that control programmes should consider distributing PBO-LLINs at more frequent intervals than three years and prioritise regions where traditional housing is common. Additionally, the contrasting performance of the same Olyset Plus nets in the WHO Cone assay and the WHO Wireball bioassay highlights that LLIN products with excito-repellent properties should be assessed with approaches that minimise avoidance from the net surface.

## Notes

### Competing Interest Statement

The authors have declared no competing interest.

## REFERENCES

1. Churcher TS, Lissenden N, Griffin JT, Worrall E, Ranson H. The impact of pyrethroid resistance on the efficacy and effectiveness of bednets for malaria control in Africa. Elife. 2016;5:e16090.

2. Pryce J, Richardson M, Lengeler C. Insecticide‐treated nets for preventing malaria. Cochrane Database of Systematic Reviews. 2018(11).

3. Bhatt S, Weiss D, Cameron E, Bisanzio D, Mappin B, Dalrymple U, et al. The effect of malaria control on Plasmodium falciparum in Africa between 2000 and 2015. Nature. 2015;526(7572):207–11.

4. Organization WH. Methods for maintaining coverage with long-lasting insecticidal nets (LLIN): Vector Control Technical Expert Group Report to MPAC. 2013.

5. Organization WH. Estimating functional survival of long-lasting insecticidal nets from field data. Vector Control Technical Expert Group Report to MPAC September 2013. 2016.

6. Gnanguenon V, Azondekon R, Oke-Agbo F, Beach R, Akogbeto M. Durability assessment results suggest a serviceable life of two, rather than three, years for the current long-lasting insecticidal (mosquito) net (LLIN) intervention in Benin. BMC infectious diseases. 2014;14(1):1–10.

7. Toé KH, Mechan F, Tangena J-AA, Morris M, Solino J, Tchicaya EF, et al. Assessing the impact of the addition of pyriproxyfen on the durability of permethrin-treated bed nets in Burkina Faso: a compound-randomized controlled trial. Malaria journal. 2019;18(1):1–16.

8. Lorenz LM, Bradley J, Yukich J, Massue DJ, Mageni Mboma Z, Pigeon O, et al. Comparative functional survival and equivalent annual cost of 3 long-lasting insecticidal net (LLIN) products in Tanzania: A randomised trial with 3-year follow up. PLoS medicine. 2020;17(9):e1003248.

9. Organization WH. Report of the twentieth WHOPES working group meeting, WHO/HQ, Geneva, 20–24 March 2017. World Health Organization; 2017.

10. Ranson H, Lissenden N. Insecticide resistance in African Anopheles mosquitoes: a worsening situation that needs urgent action to maintain malaria control. Trends in parasitology. 2016;32(3):187–96.

11. Hemingway J, Ranson H, Magill A, Kolaczinski J, Fornadel C, Gimnig J, et al. Averting a malaria disaster: will insecticide resistance derail malaria control? The Lancet. 2016;387(10029):1785–8.

12. Irish S, N’guessan R, Boko P, Metonnou C, Odjo A, Akogbeto M, et al. Loss of protection with insecticide-treated nets against pyrethroid-resistant Culex quinquefasciatus mosquitoes once nets become holed: an experimental hut study. Parasites & vectors. 2008;1(1):1–5.

13. Asidi A, N’Guessan R, Akogbeto M, Curtis C, Rowland M. Loss of household protection from use of insecticide-treated nets against pyrethroid-resistant mosquitoes, Benin. Emerging infectious diseases. 2012;18(7):1101.

14. Strode C, Donegan S, Garner P, Enayati AA, Hemingway J. The impact of pyrethroid resistance on the efficacy of insecticide-treated bed nets against African anopheline mosquitoes: systematic review and meta-analysis. PLoS medicine. 2014;11(3):e1001619.

15. Viana M, Hughes A, Matthiopoulos J, Ranson H, Ferguson HM. Delayed mortality effects cut the malaria transmission potential of insecticide-resistant mosquitoes. Proceedings of the National Academy of Sciences. 2016;113(32):8975–80.

16. Alout H, Dabiré RK, Djogbénou LS, Abate L, Corbel V, Chandre F, et al. Interactive cost of Plasmodium infection and insecticide resistance in the malaria vector Anopheles gambiae. Scientific Reports. 2016;6(1):1–11.

17. Organization WH. WHO guidance note for estimating the longevity of long-lasting insecticidal nets in malaria control. World Health Organization; 2013.

18. Gimnig JE, Lindblade KA, Mount DL, Atieli FK, Crawford S, Wolkon A, et al. Laboratory wash resistance of long‐lasting insecticidal nets. Tropical Medicine & International Health. 2005;10(10):1022–9.

19. Organization WH. Guidelines for monitoring the durability of long-lasting insecticidal mosquito nets under operational conditions. World Health Organization; 2011. Report No.: 9241501707.

20. Organization WH. Guidelines for laboratory and field-testing of long-lasting insecticidal nets. World Health Organization; 2013. Report No.: 9241505273.

21. Lynd A, Gonahasa S, Staedke SG, Oruni A, Maiteki-Sebuguzi C, Dorsey G, et al. LLIN Evaluation in Uganda Project (LLINEUP): a cross-sectional survey of species diversity and insecticide resistance in 48 districts of Uganda. Parasites & vectors. 2019;12(1):1–10.

22. Njoroge H, van’t Hof A, Oruni A, Pipini D, Nagi SC, Lynd A, et al. Identification of a rapidly-spreading triple mutant for high-level metabolic insecticide resistance in Anopheles gambiae provides a real-time molecular diagnostic for anti-malarial intervention deployment. bioRxiv. 2021.

23. Staedke SG, Kamya MR, Dorsey G, Maiteki-Sebuguzi C, Gonahasa S, Yeka A, et al. LLIN Evaluation in Uganda Project (LLINEUP)–Impact of long-lasting insecticidal nets with, and without, piperonyl butoxide on malaria indicators in Uganda: study protocol for a cluster-randomised trial. Trials. 2019;20(1):1–13.

24. Staedke SG, Gonahasa S, Dorsey G, Kamya MR, Maiteki-Sebuguzi C, Lynd A, et al. Effect of long-lasting insecticidal nets with and without piperonyl butoxide on malaria indicators in Uganda (LLINEUP): a pragmatic, cluster-randomised trial embedded in a national LLIN distribution campaign. The Lancet. 2020;395(10232):1292–303.

25. Gleave K, Lissenden N, Chaplin M, Choi L, Ranson H. Piperonyl butoxide (PBO) combined with pyrethroids in insecticide‐treated nets to prevent malaria in Africa. Cochrane Database of Systematic Reviews. 2021(5).

26. Lorenz LM, Overgaard HJ, Massue DJ, Mageni ZD, Bradley J, Moore JD, et al. Investigating mosquito net durability for malaria control in Tanzania-attrition, bioefficacy, chemistry, degradation and insecticide resistance (ABCDR): study protocol. BMC public health. 2014;14(1):1–11.

27. Sutcliffe JF, Yin S. Behavioural responses of females of two anopheline mosquito species to human-occupied, insecticide-treated and untreated bed nets. Malaria journal. 2014;13(1):1–19.

28. Sutcliffe J, Ji X, Yin S. How many holes is too many? A prototype tool for estimating mosquito entry risk into damaged bed nets. Malaria journal. 2017;16(1):1–21.

29. Ngufor C, Fagbohoun J, Agbevo A, Ismail H, Challenger JD, Churcher TS, et al. Comparative efficacy of two pyrethroid-piperonyl butoxide nets (Olyset Plus and PermaNet 3.0) against pyrethroid resistant malaria vectors: a non-inferiority assessment. Malaria journal. 2022;21(1):1–13.

30. Owusu HF, Müller P. How important is the angle of tilt in the WHO cone bioassay? Malaria journal. 2016;15(1):1–10.

31. Organization WH. Guidelines for testing mosquito adulticides for indoor residual spraying and treatment of mosquito nets. World Health Organization; 2006.

32. Okumu FO, Chipwaza B, Madumla EP, Mbeyela E, Lingamba G, Moore J, et al. Implications of bio-efficacy and persistence of insecticides when indoor residual spraying and long-lasting insecticide nets are combined for malaria prevention. Malaria journal. 2012;11(1):1–13.

33. Randriamaherijaona S, Briët OJ, Boyer S, Bouraima A, N’Guessan R, Rogier C, et al. Do holes in long-lasting insecticidal nets compromise their efficacy against pyrethroid resistant Anopheles gambiae and Culex quinquefasciatus? Results from a release–recapture study in experimental huts. Malaria journal. 2015;14(1):1–22.

34. Bertozzi-Villa A, Bever CA, Koenker H, Weiss DJ, Vargas-Ruiz C, Nandi AK, et al. Maps and metrics of insecticide-treated net access, use, and nets-per-capita in Africa from 2000–2020. Nature communications. 2021;12(1):1–12.

35. Koenker H, Kilian A, de Beyl CZ, Onyefunafoa EO, Selby RA, Abeku T, et al. What happens to lost nets: a multi-country analysis of reasons for LLIN attrition using 14 household surveys in four countries. Malaria journal. 2014;13(1):1–10.

36. Lynd A, McCall PJ. Clustering of host-seeking activity of Anopheles gambiae mosquitoes at the top surface of a human-baited bed net. Malaria journal. 2013;12(1):1–8.

37. Parker JE, Angarita-Jaimes N, Abe M, Towers CE, Towers D, McCall PJ. Infrared video tracking of Anopheles gambiae at insecticide-treated bed nets reveals rapid decisive impact after brief localised net contact. Scientific reports. 2015;5(1):1–14.

