## Supplemental material for "LLIN Evaluation in Uganda Project (LLINEUP) – The durability of long-lasting insecticidal nets treated with and without piperonyl butoxide (PBO) in Uganda"

**Supplementary material**

**Supplementary Table 1.** Sample size of nets used for each outcome**.**

| **Outcome** | **Timepoint** | **PermaNet** **2.0**  **(n)** | **PermaNet** **3.0**  **(n)** | **Olyset  Net**  **(n)** | **Olyset  Plus**  (n) | **Total** |
| --- | --- | --- | --- | --- | --- | --- |
| **Physical integrity** | 12 Month | 98 | 100 | 100 | 97 | **395** |
|  | 25 Month | 97 | 100 | 99 | 100 | **396** |
| **Chemical integrity (HPLC)** | Baseline | 5 | 5 | 5 | 5 | **20** |
|  | 12 Month | 38 | 35 | 34 | 31 | **138** |
|  | 25 Month | 29 | 30 | 30 | 30 | **119** |
| **WHO Cone assay** | Baseline | 5 | 5 | 5 | 5 | **20** |
|  | 12 Month | 7 | 7 | 7 | 7 | **28** |
|  | 25 Month | 7 | 7 | 7 | 7 | **28** |
|  | Baseline | 5 | 5 | 5 | 5 | **20** |
| **WHO Wireball assay** | 12 Month | 5 | 5 | 5 | 5 | **20** |
|  | 25 Month | 5 | 5 | 5 | 5 | **20** |

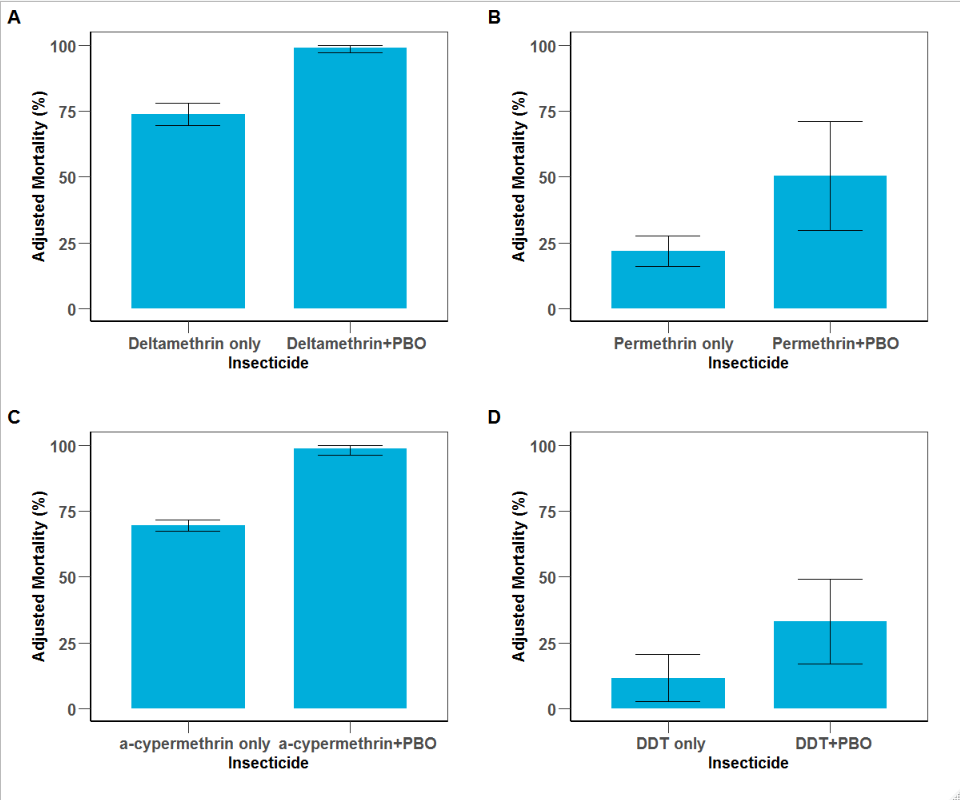
**Supplementary Figure 1.** Adjusted mortality of *An. gambiae* strain ‘Busia’ after 60 minute WHO tube exposure to **(A)** 0.05% deltamethrin, **(B)** 0.75% permethrin, **(C)** 0.05% alpha-cypermethrin, **(D)** 4% DDT.

**Supplementary Table 2** Percentage of sampled nets of each LLIN product in each pHI category at 12 and 25 months post distribution.

| **Timepoint** | **LLIN Product** | **pHI category** | | |
| --- | --- | --- | --- | --- |
|  |  | **‘Good’** | **‘Damaged’** | **‘Too torn’** |
| **12 months** | **Olyset Net** | 63.0 | 26.9 | 11.0 |
|  | **Olyset Plus** | 61.86 | 24.74 | 13.40 |
|  | **PermaNet 2.0** | 71.43 | 24.49 | 4.08 |
|  | **PermaNet 3.0** | 67.00 | 26.00 | 7.00 |
| **25 months** | **Olyset Net** | 39.39 | 44.44 | 16.16 |
|  | **Olyset Plus** | 48.00 | 30.00 | 22.00 |
|  | **PermaNet 2.0** | 55.10 | 34.69 | 10.20 |
|  | **PermaNet 3.0** | 49.00 | 36.00 | 15.00 |

**Supplementary Table 3**

| **Outcome** | **Variable** | **Odds ratio** | **95% CI** | **p** |
| --- | --- | --- | --- | --- |
| ‘too torn’ | Timepoint  (25m) | 2.017 | 1.276-3.238 | <0.001 |
|  | Housetype (Traditional) | 3.351 | 1.838-6.002 | 0.003 |
| Any holes | Timepoint  (25m) | 1.821 | 1.289-2.571 | <0.001 |

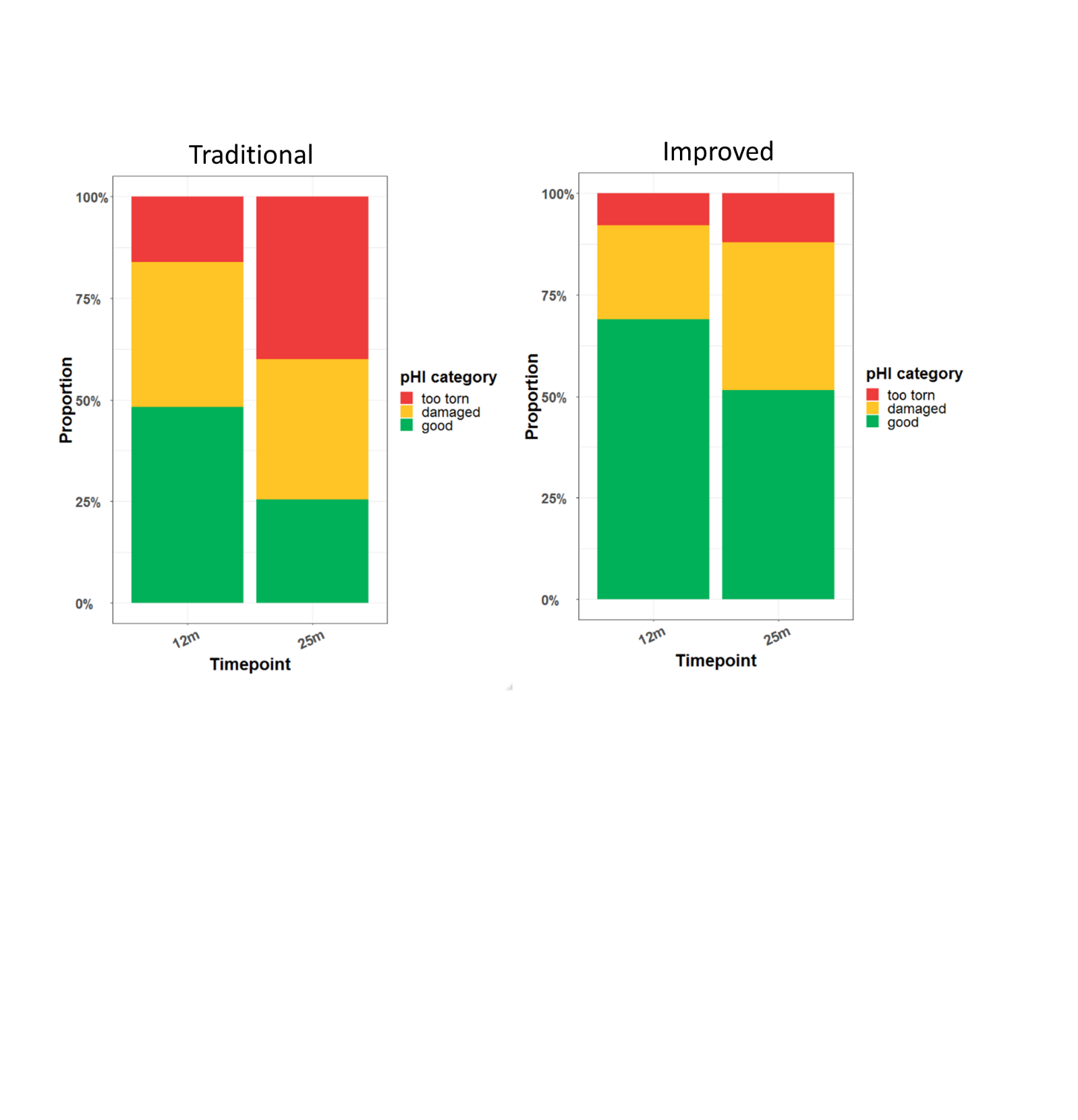
**Supplementary Figure 2.** Proportion of nets from traditional and improved housing in each pHI category at 12 and 25 months post distribution.
